# Pair housing makes calves more optimistic

**DOI:** 10.1101/691691

**Authors:** Katarína Bučková, Marek Špinka, Sara Hintze

## Abstract

Individual housing of dairy calves is common farm practice, but has negative effects on calf welfare. A compromise between practice and welfare may be housing calves in pairs. We compared learning performances and affective states as assessed in a judgement bias task of individually housed and pair-housed calves. Twenty-two calves from each housing treatment were trained on a spatial Go/No-go task with active trial initiation to discriminate between the location of a teat-bucket signalling either reward (positive location) or non-reward (negative location). We compared the number of trials to learn the operant task (OT) for the trial initiation and to finish the subsequent discrimination task (DT). Ten pair-housed and ten individually housed calves were then tested for their responses to ambiguous stimuli positioned in-between the positive and negative locations. Housing did not affect learning speed (OT: F_1,34_ = 0.42, P = 0.52; DT: F_1,34_ = 0.25, P = 0.62), but pair-housed calves responded more positively to ambiguous cues than individually housed calves (χ^2^_1_ = 6.76, P = 0.009), indicating more positive affective states. This is the first study to demonstrate that pair housing improves the affective aspect of calf welfare when compared to individual housing.

## Introduction

Most farm animals, including dairy cattle, are social species with a strong tendency to form groups^1^. Under free-ranging conditions, cows separate themselves from the herd shortly before parturition to give birth in a hidden place^2^. With two to three days of age, calves are introduced to the herd^3^, and at the age of three weeks, they spend already most of their time in the company of other calves^4^.

Dairy calves reared for commercial purposes, on the contrary, are commonly separated from their mothers within 24 hours after birth^5^. In Europe, 60 % of dairy calves are housed individually during their first eight weeks of life^6^. In some European countries, individual housing of dairy calves in early ontogeny prevails even more. For example, in Czech Republic, dairy calves are housed individually on 97 % of the farms^7^. The main declared aims of individual housing are to minimise the spread of diseases and to reduce cross-sucking and competition over milk^8^. Moreover, individually housed calves are easier to observe and treat^9^. However, in comparison with housing in groups, individual housing provides calves with less space to move^10^, fewer opportunities to play^11,12^ and to learn social skills^8,10,13^. Individually housed calves have also more difficulties in coping with novel situations^8,11^. Furthermore, several studies found that calves housed individually take in less solid feed^8^ and grow more slowly than calves housed in groups^8,12^.

Compared to group housing, pair housing of calves may not provide the full benefits of group housing, but may be more likely implemented by farmers as it facilitates observation and manipulation of animals and may evoke less concern over spread of pathogens, occurrence of cross-sucking or milk competition. Therefore, pair housing may be a good compromise between group housing and individual housing in terms of calf welfare and farm practice. Studies across species, including birds (e.g. parrots^14^) and mammals (e.g. rodents^15^, horses^16,17^, primates^18^ or dogs^19^) show that housing in pairs provides animals with welfare benefits compared to individual housing, including a decrease in abnormal and stress-related behaviours, although it is not clear whether these benefits match those provided by group housing. Findings in calves also suggest that pair housing has the potential to improve calf welfare. Calves housed in pairs have been shown to take in more solid feed and to gain more weight^20^. Moreover, changes in behaviour (e.g. reduced weaning stress^6,21^ or less reactivity to novelty^22^) as well as in learning abilities (better performance in reversal learning tasks^22,23^) have been identified between pair-housed and individually housed calves.

Improved learning abilities of pair-housed dairy calves may help the animals to cope with changes in their routine management later in life when dairy cattle are faced with many challenges, including interaction with new equipment and changes in social structure, feeding environment or staff. More flexible individuals may adapt more quickly to these challenges, which may improve their welfare and facilitate the work of stockpersons^22^. To our knowledge, only two studies have compared the learning abilities of individually housed and pair-housed calves so far. Both Gaillard et al.^22^ and Meagher et al.^23^ found that individually and pair-housed calves did not differ in learning to discriminate between two colours, but that pair-housed calves performed better in learning a reversal task. Moreover, the study by Meagher et al. revealed that the majority of individually housed calves did not learn the reversal task at all, even when provided with twice as many sessions as required by an average pair-housed calf^23^.

Taken together, rearing calves in pairs improves several aspects of their welfare, potentially leading to more positive affective states in pair-housed compared to individually housed calves. However, so far no study has investigated how individual housing and pair housing influence calves’ affective states. One promising approach to assess the valence of affective states in animals (i.e. whether an animal experiences its situation as pleasant or unpleasant) is to record their cognitive biases. From humans we know that affective states influence how a person sees the world^24^; they can thus bias different cognitive processes, including memory^25^, attention^26^ and judgement^27^. Most research on cognitive biases in non-human animals has focused on judgement biases as assessed in an individual’s responses to ambiguous cues^28^. The Go/No-go task, the most commonly used judgement bias task in animals, was initially validated in rats^29^ and consists of two stages: In the first stage, animals are trained on an discrimination task to show a Go response (i.e. approach of a goal to receive a reward) when exposed to one cue (positive trial), and to show a No-go response (i.e. no approach of the goal since there is no reward) when exposed to a different cue (negative trial). In the second stage, animals are presented with intermediate and thus ambiguous cues interspersed among the positive and negative reference cues. Animals in a positive affective state are more likely to respond to these ambiguous cues as if they predicted the positive cue and thus to show a Go response (indicating expectation of the positive outcome and thus interpreted as an ‘optimistic’ response) whereas animals in a more negative affective state are more likely to show a No-go response (indicating expectation of the negative outcome and thus interpreted as a ‘pessimistic’ response). The ratio of optimistic to pessimistic responses to the ambiguous cues is then interpreted in terms of positive and negative affective valence^28^.

Judgement bias tasks have been used across many species, including laboratory rodents (e.g. rats^29^ and mice^30^), companion animals (e.g. dogs^31^ and horses^32^) and farm animals (e.g. pigs^33^, laying hens^34^ and goats^35^). Three studies assessed judgement biases in dairy calves^36–38^. Neave et al.^36^ found that calves show a more negative judgement bias after being subjected to the painful procedure of hot-iron disbudding and Daros et al.^37^ demonstrated a similar negative judgement after separating calves from their mothers in spite of the fact that before separation calves did not have access to the cows’ udder. Whereas these two studies used task designs based on visual discrimination between two colours^36,37^, Lecorps et al.^38^ used a spatial task design and showed that more fearful calves made more negative judgements than less fearful individuals. Judgement bias tasks have not yet been used to assess calves’ affective states with respect to individual or pair housing. Only in canary birds, it has been shown that pair-housed individuals made more optimistic responses in ambiguous trials than their individually housed conspecifics^39^.

In the present study, we aimed to assess affective states of individually and pair-housed calves on a spatial Go/No-go task design with active trial initiation. To this end, we adapted to dairy calves the task design developed and successfully used in mice, rats and horses by Hintze et al.^40^. Training animals to actively initiate each trial has the advantage that they get the opportunity to skip waiting time when opting for a No-go response and to proceed immediately to the next trial, thus reducing frustration and maximising overall training speed by giving control to the animals. Besides assessing calves’ affective states, we aimed to compare their learning performances during both learning phases: First the initial operant learning phase where calves learned to initiate each trial followed by the discrimination learning phase where calves were trained to discriminate between positive and negative cues and to respond accordingly (i.e. with a Go or a No-go response, respectively).

We hypothesised that compared to individually housed calves, pair-housed calves will (1) learn more quickly both the initial operant task of active trial initiation and the subsequent discrimination between the positive and the negative cues in a spatial Go/No-go task and (2) show a higher proportion of Go responses to ambiguous cues, thus indicating more positive affective states.

## Animals, Material and Methods

### Animals and housing treatments

Our experiment was carried out at the experimental farm Netluky of the Institute of Animal Science in Prague, Czech Republic, from June 2017 to August 2018. In total, we used 66 Holstein Friesian female calves. Calves were separated from their mothers within 12 hours after birth and kept individually until they entered our experiment with an age of four to eleven days. Twenty-two calves were assigned to individual housing (IND) and 44 calves to pair housing (PAIR). Allocation to the treatments was balanced across age and weight. All IND calves and 22 PAIR calves (one focal calf per pair, chosen randomly) were trained and, if they reached learning criterion in a given time (see below), they were tested on a spatial Go/No-go task with trial initiation.

IND calves were kept in standard single pens (1.4 × 2.6 m; Supplementary Figure S1) bedded with straw. They could have visual and tactile contact with calves in neighbouring pens. PAIR calves were housed in double-sized pens (2.8 × 2.6 m; Supplementary Figure S2). All calves had free access to water and calf starter diet as soon as they were separated from their mothers. They were checked daily for their health by the researchers and farm staff and, if needed, were treated by a veterinarian.

In the first week of the experiment, calves were fed 7 l of milk per day (3.5 l at 6 a.m. and 3.5 l at 6 p.m.) via teat-buckets. In the second week, the amount of milk was increased to 8 l per day (2 l at 6 a.m., 4 l during training/testing and 2 l at 6 p.m.). From the third week onwards, calves had free access to hay and were fed 10 l of milk per day divided into three meals: 2.5, 3 or 3.5 l at 6 a.m., with the exact amount depending on how early in the day the training started, 4 l during training/testing, and the remainder to 10 l at 6 p.m.

### Experimental arena and procedures

#### Experimental arena

Calves were trained and tested in a rectangular arena (2.8 m × 4.2 m; Figure 1) located in the calf barn where all calves were housed. Five goal-holes (of round shape, 6.5 cm in diameter) were drilled in one of the arena walls at a height of 0.6 m and spaced equidistantly (0.3 m) from each other (Supplementary Figure S3). Above each goal-hole, a bucket holder was mounted on the outside of the wall (Supplementary Figure S4), allowing the researcher to place a teat-bucket in a way that calves inside the arena could only see the red teat, but not the whole teat-bucket, which was mounted outside. Opposite to the five goal-holes, a round metal object (25.5 cm in diameter) functioning as trial initiator was suspended from the ceiling; via a pulley it could be lifted and lowered by the researcher positioned behind the wall with the five goal-holes.

**Figure 1.**
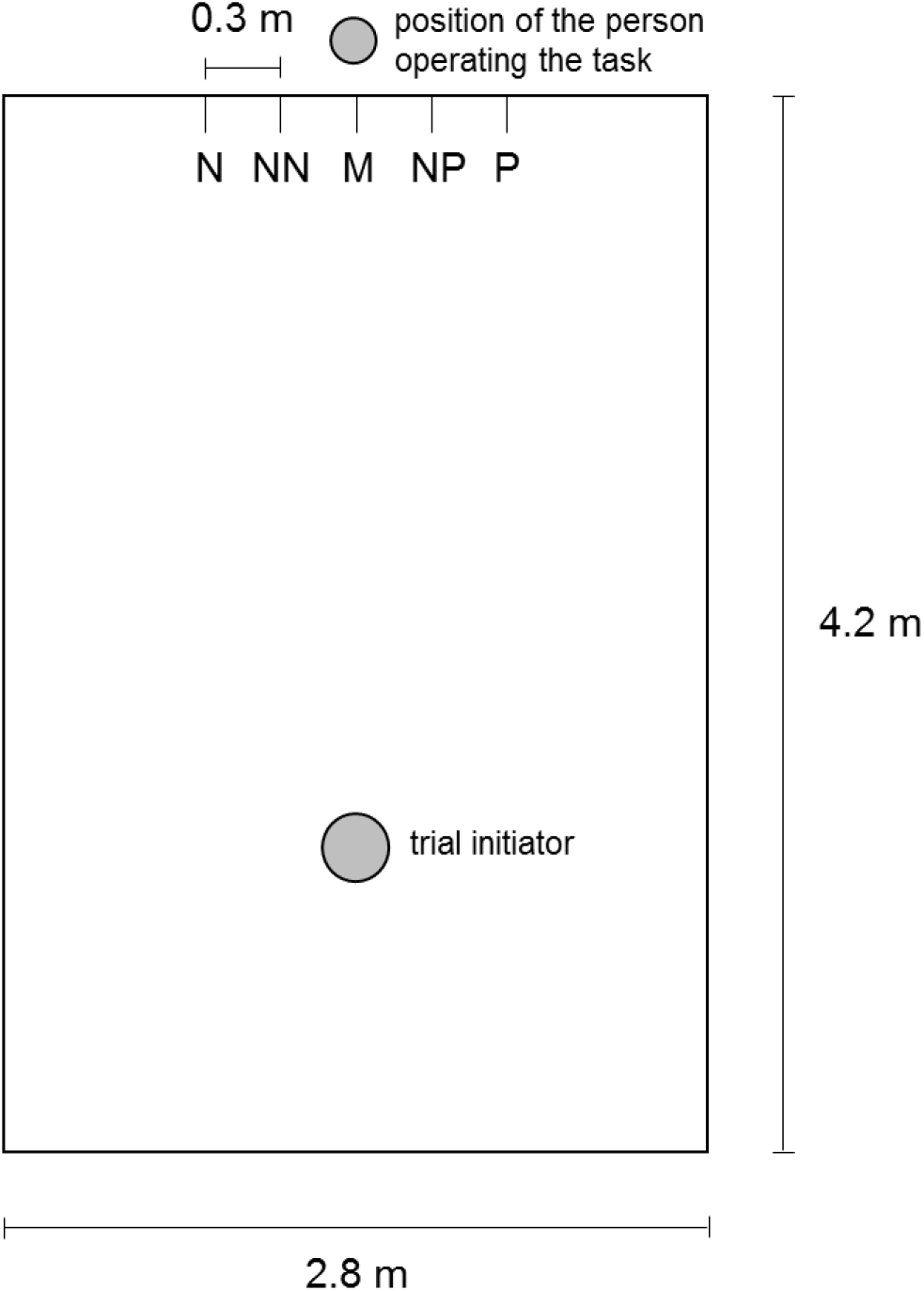
Schematic overview of the experimental arena. Overview of the experimental arena with the trial-initiator on one side and the five goal-holes on the opposite side; Positive (P), Negative (N) and the three ambiguous goal-holes (Near Positive - NP, Middle - M and Near Negative - NN).

#### Experimental procedures

Calves were trained and tested once a day on five consecutive days per week for a maximum of 30 training days. Training was then stopped because calves were disbudded, a procedure which would have interfered with training and testing. All calves were trained and tested by the same researcher and just in case of urgency by another trained researcher, who was familiar with the calves. Training consisted of two phases: The operant learning phase, in which calves learned to initiate trials and the discrimination learning phase, in which they were trained to discriminate between a positive and a negative cue and to respond accordingly by showing a Go or a No-go response, respectively. If successful, training was followed by testing calves in the judgement bias task.

##### Habituation

Before the start of training, each calf was individually habituated to the experimental arena by allowing it free exploration for 15 minutes.

##### Operant learning for trial initiation

In a stepwise process, calves were trained to initiate each trial by touching the trial initiator with the muzzle and to subsequently approach the positive location (P) to receive 0.25 l of milk as a reward. Each session consisted of 16 trials and the position of P (right/left goal-hole) was balanced across calves from both treatments.

First, calves were trained to touch the trial initiator held by the researcher very close to P. The distance between trial initiator and P was then gradually increased up to the final distance of 3 m (the different sub-phases are described in the Supplementary Material). At this stage, the trial initiator was suspended from the ceiling and controlled by the researcher behind the wall with the five goal-holes. When a calf touched the trial initiator, it was lifted, indicating that the calf had performed the correct behaviour and the teat-bucket was placed at P. If the calf then touched the teat of the bucket, milk was poured into the bucket. While the calf was sucking, the trial initiator was lowered and thus made available to be touched again. When the calf turned around to touch the trial initiator again, the bucket was removed. To reach the learning criterion for the operant learning phase, calves were allowed a maximum of one mistake per session. A mistake was defined as not touching the trial initiator within 90 s after the calf had finished drinking milk. If a calf did not touch the trial initiator within 120 s or did not start drinking the milk within 90 s after having touched the trial initiator, the session was terminated.

##### Discrimination learning

After having reached the learning criterion for the operant learning, calves were trained to show Go responses in positive and No-go responses in negative trials, the latter being now introduced. A Go response was defined as the calf touching the teat of the bucket. A No-go response was defined as initiation of a new trial by touching the trial initiator within 90 s after cue presentation. The procedure during positive trials remained the same as during the operant learning. In negative trials, the bucket was placed into the bucket holder at the location opposite of P (negative location, N). When the calf touched the teat of the bucket at N, it did not get any milk, but was allowed to initiate the next trial. In the first session, 16 positive and 8 negative trials were presented. From the second session onwards, sessions consisted of 16 positive and 16 negative trials. Trials were presented in pseudorandom order with no more than two consecutive positive or negative trials, but with the first and the last trial always being positive. To fulfil the discrimination criterion, calves had to show at least 13 Go responses out of the 16 positive and 13 No-go responses out of the 16 negative trials in two consecutive sessions. If a calf neither started to drink the milk (in a positive trial) nor touched the trial initiator within 90 s, the session was terminated.

##### Judgement bias testing

Once a calf had reached the discrimination criterion, it was tested in up to four test sessions (fewer if a calf got sick or 30 days for training and testing had already expired). Each test session consisted of 16 positive, 16 negative and three ambiguous trials, with all three ambiguous cue locations (Near Positive - NP, Middle - M, Near Negative - NN) being presented once per test session. The order of positive and negative trials corresponded to the same rules as during discrimination learning. Ambiguous trials were balanced equally often after positive and negative trials across the four test sessions, and the order of each ambiguous cue within a test session was balanced across sessions. Go and No-go responses were defined as during discrimination learning. Go responses in positive trials were rewarded, Go responses in negative trials were not rewarded and Go responses in ambiguous trials were rewarded to reduce potential effects of surprising non-reward^40–42^. According to the principles of the judgement bias task^29^, Go responses in ambiguous trials were interpreted as ‘optimistic’ responses, whereas No-go responses were interpreted as ‘pessimistic’ responses. If a calf did neither start drinking the milk (in a positive trial) nor touched the trial initiator and thus re-initiated in a negative trial within 90 s after cue presentation, the test session was terminated. If a calf did not initiate all 35 trials or made more than four mistakes in the positive trials and/or more than four mistakes in the negative trials, it was tested in up to two additional test sessions.

##### Exclusion criteria

During the course of training, calves were excluded for various reasons. A calf was discarded for reasons of ‘Poor Health’ (PH) if 1) it had diarrhoea for more than 12 consecutive days or was generally sick for more than five consecutive days or 2) its non-focal social partner was in a generally bad condition for at least five consecutive days, which might have affected the respective focal calf. A calf was excluded because of ‘Low Motivation’ (LM) if it did not initiate the first trial of a session within 30, 60 or 90 s (depending on the sub-phase of operant learning as described in the Supplementary Material) in more than two consecutive sessions. If a calf did not meet the learning criterion for the discrimination learning task within 30 days of training, it was classified as ‘Not Learned’ (NL).

#### Ethical considerations

The experimental protocol was approved by the Institutional Animal Care and Use Committee of the Institute of Animal Science.

#### Statistical analyses

##### Learning performance

Training duration and attrition rate for training are presented descriptively per housing treatment. Calves’ learning performances were compared based on the number of trials (mean ± standard deviation; SD) needed to fulfil the operant learning criterion (IND: = 20, PAIR: n = 17) and to reach the operant plus discrimination learning criterion (IND: n = 10, PAIR: n = 11). The number of trials needed for the discrimination learning alone could not be analysed separately because the length of the operant learning affected the maximum number of sessions a calf could be trained for in the discrimination learning phase since training was stopped after 30 training days; the more sessions it needed to fulfil the operant learning criterion, the fewer days were left to reach the discrimination learning criterion. To investigate the effect of individual versus pair housing on calves’ learning performances, linear models with housing treatment (IND, PAIR) as fixed effect, age at the start of learning (range between 11 and 18 days) as covariate and number of trials (log-transformed, if needed to achieve quasi normal distribution) as the dependent variable were run. Models were checked for normality of residuals and for homoscedasticity through plotting the residuals against the fixed effect and the covariate. Data were analysed in SAS (version 9.4).

##### Judgement bias testing

A stable performance in positive and negative trials during test sessions is a prerequisite for a valid interpretation of the animals’ responses in ambiguous trials^43^. To this end, test sessions in which calves showed fewer than 13 Go responses in positive trials and fewer than 13 No-go responses in negative trials were excluded from further analyses (comparable to how it was done by Hintze et al.^40^). To investigate the effect of individual (n = 10) versus pair housing (n = 10) on calves’ decisions in the ambiguous trials (binary dependent variable: Go response = 1, No-go response = 0), generalized mixed-effects models were run (function glmer of the package lme4; ‘family’: binomial, including the ‘logit’ link function^44^). Fixed effects were housing treatment (IND, PAIR) and age of the calves on the first day of test (range between 26 and 62 days). Random effects were ambiguous trial type (NP, M, NN) nested in test session (1 - 4) nested in calf. To analyse the effect of housing treatment on calves’ responses in positive and negative trials, generalized mixed-effects models with housing treatment as fixed effect and session nested in calf as random effects were run. Data were analysed in R (version 3.5.2).

## Results

### Learning performance

#### Training duration

IND calves needed between 95 and 271 trials (170 ± 52.4; i.e. about 10.2 ± 3.2 sessions) and PAIR calves needed between 96 and 321 trials (185 ± 67.2, i.e. about 11.5 ± 4.2 sessions) to learn the operant task for trial initiation. In total (operant learning plus discrimination learning), it took IND calves between 408 and 678 trials (534 ± 90.7, i.e. about 22.1 ± 3.2 sessions) and PAIR calves between 317 and 645 trials (516 ± 110, i.e. 22.4 ± 4.1 sessions) to reach the discrimination learning criterion. Housing treatment did not affect calves’ learning performances, neither in the operant learning phase (F_1,34_ = 0.42, P = 0.52) nor in the whole learning process until the end of the discrimination learning (F_1,18_ = 0.25, P = 0.62, Figure 2). Calves’ learning performances were not affected by their age at the start of training (operant learning: F_1,34_ = 0.33, P = 0.57; operant plus discrimination learning: F_1,18_ = 0.54, P = 0.47).

**Figure 2.**
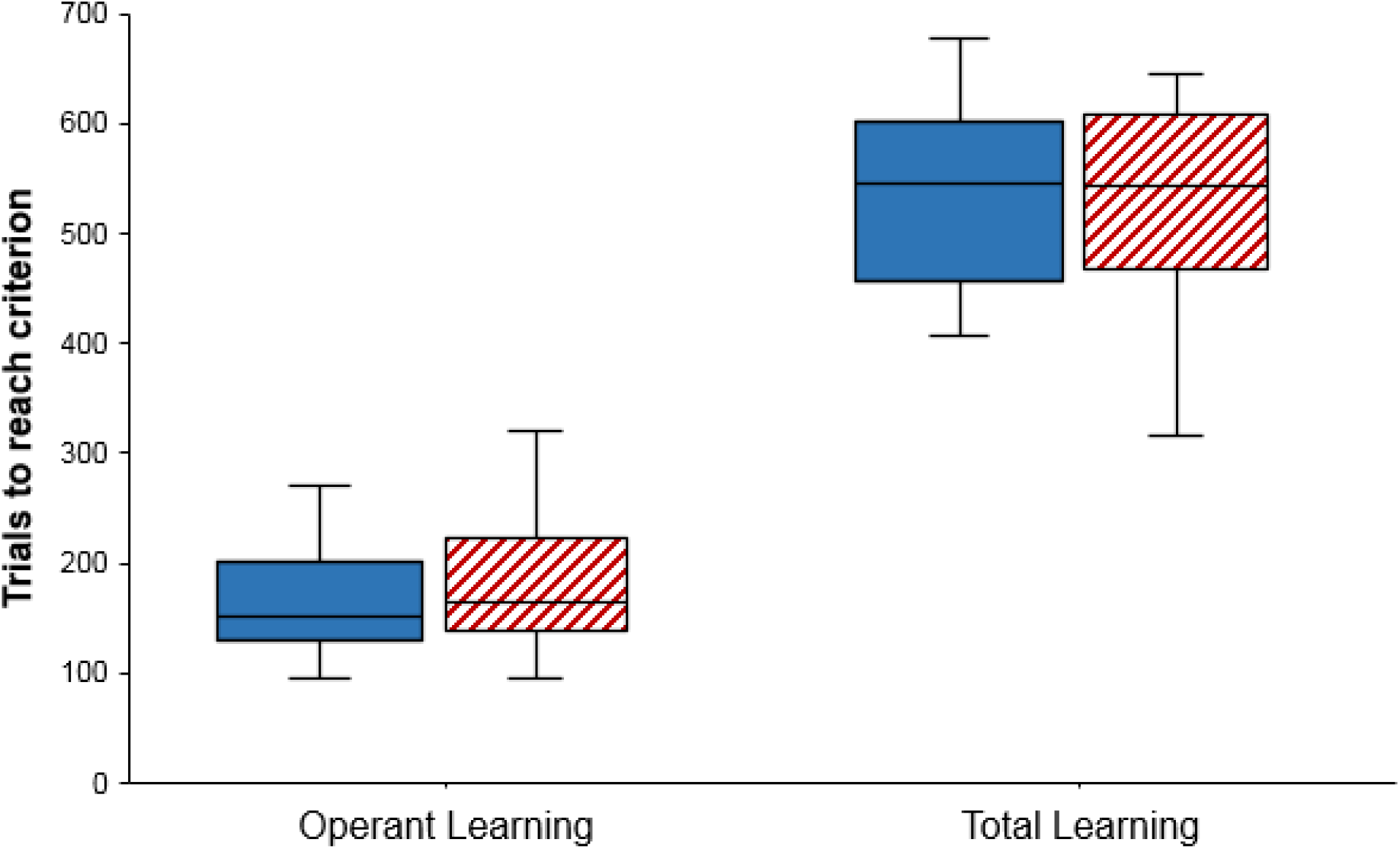
Number of trials needed to reach the criterion for operant learning and for operant plus discrimination learning (total learning). The boxplots depict median, interquartile range and data range. Open blue boxes: IND calves (n = 20 for operant learning and n = 10 for total learning). Hatched red boxes: PAIR calves (n = 17 for operant learning and n = 11 for total learning).

#### Attrition rate

The number of discarded calves per training phase and housing treatment and the reasons for exclusion are given in Table 1.

**Table 1.** Number of discarded calves per training phase and housing treatment, and reasons for attrition.

| Phase | Total number of discarded calves | Number of discarded IND calves |  | Number of discarded PAIR calves |  |
| --- | --- | --- | --- | --- | --- |
|  |  | Number | Reason | Number | Reason |
| Operant learning | 7 | 2 | PH: n = 2 | 5 | PH: n = 1 |
|  |  |  | LM: n = 0 |  | LM: n = 4 |
| Discrimination learning | 16 | 10 | PH: n = 0 | 6 | PH: n = 0 |
|  |  |  | LM: n = 1 |  | LM: n = 0 |
|  |  |  | NL: n = 9 |  | NL: n = 6 |

PH: 'Poor Health', LM: 'Low Motivation', NL: 'Not Learned' (definitions are given in the section *Exclusion criteria*).

### Judgement bias testing

Housing treatment affected calves’ responses in ambiguous trials (χ^2^_1_ = 6.76, P = 0.009; Figure 3) with PAIR calves showing Go responses in 51.4 % (± SE 29.7) of all ambiguous trials, while IND calves did so in 24.7 % (± SE 14.3) of the trials (Odds Ratio = 2.50). Age of the calves at the beginning of testing did not affect responses to ambiguous cues (χ^2^_1_ = 0.16, P = 0.69; Odds Ratio = 1.01). In accordance with the assumptions underlying the principle of the judgement bias task, PAIR calves did not differ from IND calves in their responses in positive (χ^2^_1_ = 0.42, P = 0.52) or negative trials (χ^2^_1_ = 1.16, P = 0.28).

**Figure 3.**
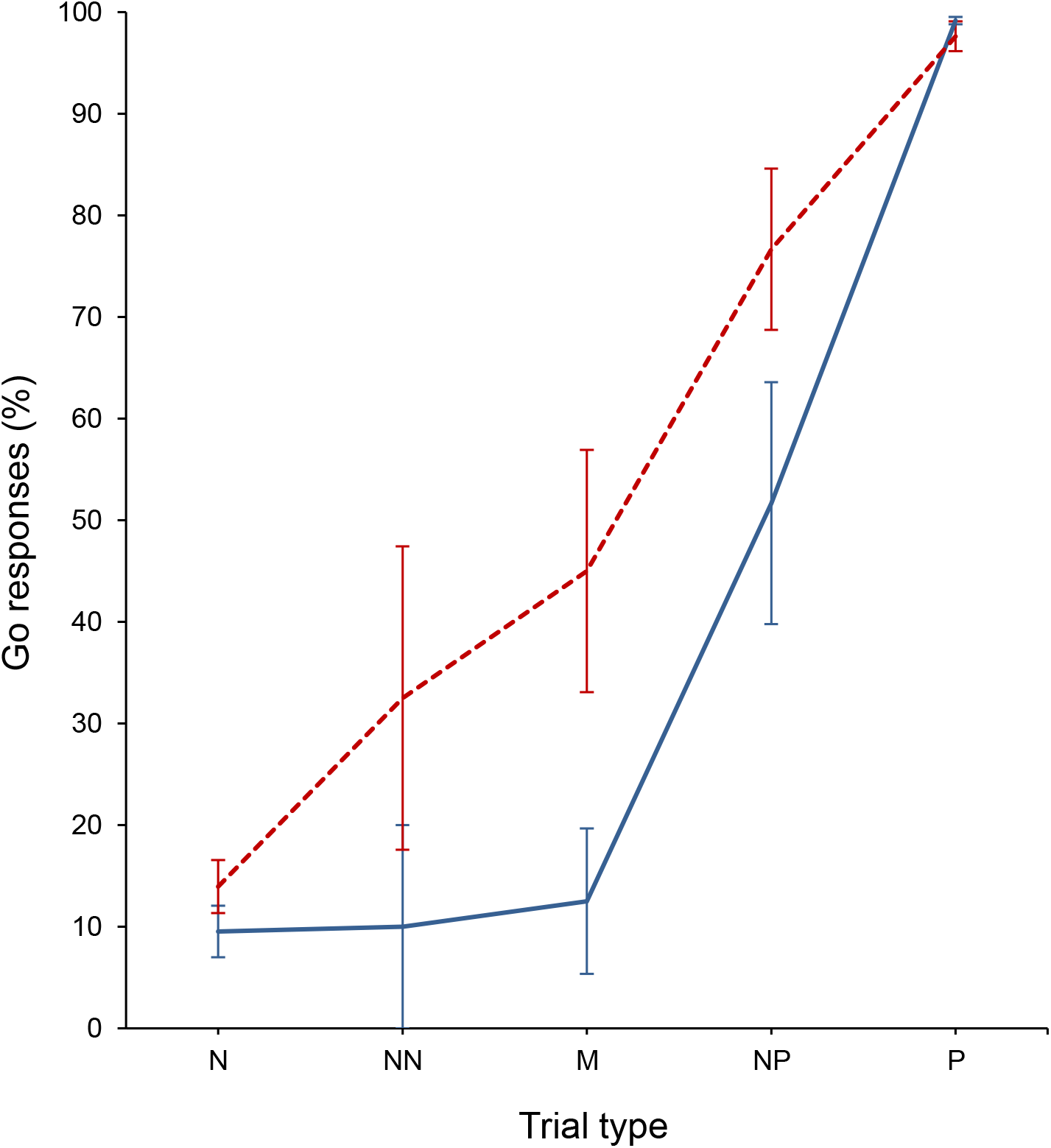
Go responses of calves in positive (P), negative (N) and the three ambiguous trial types (NP, M, NN) shown as mean ± SE. Solid blue line: IND calves (n = 10). Dashed red line: PAIR calves (n = 10).

## Discussion

Our study aimed to compare calves’ learning performances and affective states as assessed on a judgement bias task with respect to pair (PAIR) and individual (IND) housing. We found that housing treatments did not affect learning performance, neither in the initial operant task for active trial initiation nor in the total learning process, but that PAIR calves showed more Go responses in ambiguous trials, indicating that they were in a more positive affective state than IND calves.

Our results do not support the notion that having social company generally enhances the learning performance of mammals^45^, but they are consistent with the results by Gaillard et al.^22^ and Meagher et al.^23^, who did not find an effect of individual versus social housing (pair and group housing, respectively) on calves’ discrimination learning speed. However, other types of learning might be affected by social as opposed to individual housing of calves. For instance, the two cited studies^22,23^ found that pair-housed and group-housed calves were faster in reversal learning than their individually housed conspecifics. Thus, it is possible that social housing does not enhance performance in operant or discrimination tasks, but that it might help to adjust to rapidly changing environments where reversals of already learned associations and skills are needed. Furthermore, many social learning skills, e.g. individual recognition or learning through local enhancement, have not yet been studied in calves reared in different social environments.

Since PAIR and IND calves did not differ in learning speed, data on training duration from both housing treatments can be pooled for comparison with other studies. Calves in our study took in average 177 trials (around 11.1 sessions) to successfully learn the operant task for active trial initiation. To our knowledge, active trial initiation has only been implemented in three judgement bias studies so far. Neave et al.^36^ used it in dairy calves, but only introduced it at a later stage, which is why their results are not comparable to ours. Rats in the study by Jones et al.^46^ were trained to initiate trials by performing a nose-poke, but here again the training schedule differed from ours, because rats were not only trained to initiate each trial, but also to stay in this nose-poking position when a positive tone was played in order to receive a reward. The way we introduced the trial initiation is most similar to how Hintze et al.^40^ implemented it in their study on horses, rats and mice. All three species in their study needed on average fewer sessions (horses: 3.4 sessions, rats: 6.2 sessions, mice: 7.0 sessions) to successfully learn the task than our calves. However, it is important to stress that calves in our study were trained gradually to meet specific criteria for each of three sub-phases during the operant learning (see Supplementary Material), which is one possible explanation for the difference in training speed. Moreover, our calves were younger than animals of the three species Hintze et al. worked with, rendering a direct comparison difficult. We are positive that training calves in the operant task can be sped up in future studies, e.g. by reducing the number of sub-phases with required learning criteria, and by giving calves additional signals when they perform the correct behaviour, for instance by using a clicker as it was done by Neave et al.^36^.

Calves had a maximum of 30 training days (and thus sessions) to learn both the operant and the discrimination task; individuals that did not meet the learning criterion within that time were dropped. This restricted number of training sessions resulted in a relatively high attrition rate, with nine IND and six PAIR calves not learning the task (‘NL’). Calves meeting the learning criterion needed on average 357 trials (i.e. around 11.2 sessions) to successfully learn the spatial discrimination, which is comparable to calves in the study by Neave et al.^36^ that took on average 14.5 sessions for visual discrimination with a similar number of trials per session (average of 30.5 trials). Compared to the study by Hintze et al.^40^, calves needed fewer trials than the horses, but more trials than the rodents. However, due to the lower number of trials per session (32 in our study compared to 50 in the study by Hintze et al.^40^), session number was higher in our study, also when compared to horses. The number of trials a session can contain depends on the animals’ motivation to perform the task. Calves in the studies by Neave et al.^36^ and Daros et al^37^. were fed 0.14 l milk per positive trial (compared to 0.25 l milk in our study) and performed in on average 55 trials and 60 trials per session, respectively. Thus, dividing the total daily milk allotment into smaller amounts per trial might speed up the discrimination learning in terms of required sessions in future studies.

Building on the task design developed by Hintze et al.^40^ in other species, we here provide a first protocol for assessing dairy calves’ affective states by using a spatial Go/No-go task with active trial initiation. Using trial initiation enables animals to skip waiting time when opting for a No-go response and to proceed immediately to the next trial. The study by Hintze et al.^40^ showed that the theoretical assumptions underlying judgement bias tasks as proposed by Gygax^43^ were met with respect to internal task validity. Here again, these assumptions were fulfilled as calves in our study showed mostly No-go responses in negative trials, Go responses in positive trials and a monotonically graded response in ambiguous trials with the proportion of Go responses increasing the closer the ambiguous cue was to the positive cue (increase from NN to M to NP). Such a monotonically graded response pattern is crucial to ensure that animals judge the ambiguous cues in relation to the positive and negative reference cues, which function as anchor points. On top of these aspects of internal task validity, the two assumptions with respect to predictive validity proposed by Gygax^43^ were met in our study. First, calves from both housing treatments did not differ in their responses in positive and negative reference trials during testing. Second, calves did differ in their responses in ambiguous trials in that PAIR calves had 2.5 times higher odds to show Go responses across the three ambiguous trial types than IND calves, indicating that PAIR calves interpreted the ambiguous cues more optimistically (i.e. in expectation of a positive outcome) and that they were thus in a more positive affective state than IND calves. The direction of the treatment differences was consistent across all three ambiguous cues (i.e. PAIR calves showed more Go responses in NN, M and NP trials), indicating that our treatment effect is relatively robust compared to other studies, where animals from one treatment sometimes show a positive bias in one but a negative bias in another ambiguous trial type, rendering the findings hard to interpret^47,48^.

The specific reason why PAIR and IND calves differed in affective states could not be established from our data since the presence/absence of a social companion and absolute space allowance were confounded in our study, with PAIR calves not only having a companion but also having double the amount of space compared to IND calves (while space allowance per individual was the same). The effect of social companionship and space allowance was not disentangled since pair housing in commercial practice provides calves with both aspects, and we were specifically interested in studying whether such a change in housing practice would result in better welfare with respect to calves’ affective states. From other studies we know that calves are more motivated to get access to full contact than to head contact with a familiar conspecific^49^ and it is therefore possible that the relatively more negative affective state of IND calves was caused by a lack of full social contact with a conspecific. Moreover, the difference in affective states between PAIR and IND calves might be explained by a reduced amount and perhaps reduced quality of play behaviour. It is known that calves in individual housing play less^11^ and that the rebound in play behaviour when individually housed calves are exposed to a larger space and/or companions documents that these calves were play-deprived before^12^. Independent of whether it was one specific or a combination of different aspects leading to the differences in calves’ responses in ambiguous trials, our study shows that calves’ housing conditions during early ontogenesis play a crucial role for their welfare. Their comparably more positive affective states might enable PAIR calves to provide more effective social support to their partners. It is known that social partners decrease the effects of stressors in mammals^50–52^ and that this effect is stronger with familiar compared to unfamiliar conspecifics (e.g. in calves^53^). Thus, our study might incite a novel line of research, namely to investigate whether calves (and social animals in general) that are in a more positive affective state provide more effective social support than individuals that are in a more negative affective state.

Even though our results show that PAIR calves responded more optimistically than IND calves, we do not know whether the more positive affective states of PAIR calves are comparable to the affective states of group-housed calves, which is why a comparison between pair-housed and group-housed calves would be very valuable. Furthermore, it would be interesting to study the long-term effects of different housing conditions on the animals’ affective states, which, if longer-lasting, might help the animals to better cope with challenges and to become more resilient, potentially resulting in both better calf and later cow welfare and higher productivity.

In conclusion, we showed that housing treatments did not affect training duration to learn a spatial judgement bias task with active trial activation but that PAIR calves judged ambiguous stimuli more positively than IND calves. In addition to previous studies that focused on the effects of housing on health, performance and behavioural aspects of welfare, this is the first study demonstrating that pair housing improves the affective aspect of calf welfare, i.e. how calves feel about their situation^54^ when compared to individual housing.

## Supporting information

Supplementary material to Buckova et al_Manuscript

## Acknowledgements

We would like to thank Radka Šárová for her help with designing the experiment, Zuzana Andrejchová for her assistance with the training of the calves and Ágnes Moravcsíková for her help with preparing the data for analysis. Additionally, we would like to thank Iva Leszkowová for her help with taking care of the calves and the farm staff for their help with building our experimental arena and with taking care of the calves when necessary. This research project has been financed by grant No. MZE-RO0719 from the Czech Ministry of Agriculture.

## Author contributions

K.B. designed and ran the experiment, collected all data, prepared data for analysis, wrote the paper. M.Š. obtained funding, provided advice on the methodology of the experiment, analysed the data on learning performance, wrote the paper. S.H. provided advice on the design of the judgment bias task, analysed the data of the judgement bias task, wrote the paper.

## Competing interests

The authors declare no competing interests.

## Data availability

The datasets generated during this study are available in the Figshare repository.

## Notes

https://figshare.com/articles/Learning_performance_and_judgement_bias_testing_data_xlsx/8427113

https://figshare.com/articles/Untitled_Item/8427743

## References

1. Estevez, I., Andersen, I. L. & Nævdal, E. Group size, density and social dynamics in farm animals. Appl. Anim. Behav. Sci. 103, 185–204 (2007).

2. Lidfors, L. M., Moran, D., Jung, J., Jensen, P. & Castren, H. Behavior at calving and choice of calving place in cattle kept in different environments. Appl. Anim. Behav. Sci. 42, 11–28 (1994).

3. Vitale, A. F., Tenucci, M., Papini, M., Lovari, S. Social behaviour of the calves of semi-wild Maremma cattle, Bos primigenius taurus. Appl. Anim. Behav. Sci. 16, 217–231 (1986).

4. Sato, S., Wood-Gush, D. G. M. & Wetherill, G. Observations on creche behavior in suckler calves. Behav. Processes 15, 333–343 (1987).

5. Weary, D. M. & Chua, B. Effects of early separation on the dairy cow and calf: 1. separation at 6 h, 1 day and 4 days after birth. Appl. Anim. Behav. Sci. 69, 177–188 (2000).

6. Bolt, S. L., Boyland, N. K., Mlynski, D. T., James, R. & Croft, D. P. Pair housing of dairy calves and age at pairing: effects on weaning stress, health, production and social networks. PLoS One 12, e0166926 (2017).

7. Staněk, S., Zink, V., Doležal, O. & Štolc, L. Survey of preweaning dairy calf-rearing practices in Czech dairy herds. J. Dairy Sci. 97, 3973–3981 (2014).

8. Costa, J. H. C., von Keyserlingk, M. A. G. & Weary, D. M. Invited review: Effects of group housing of dairy calves on behavior, cognition, performance, and health. J. Dairy Sci. 99, 2453–2467 (2016).

9. Rushen, J., de Passilé, A. M., von Keyserlingk, M. A. G., Weary, D. M. Housing for unweaned calves in The welfare of cattle (ed. Phillips, C.) 187 (Springer, 2008).

10. Chua, B., Coenen, E., van Delen, J. & Weary, D. M. Effects of pair versus individual housing on the behavior and performance of dairy calves. J. Dairy Sci. 85, 360–364 (2002).

11. Jensen, M. B., Vestergaard, K. S., Krohn, C. C. & Munksgaard, L. Effect of single versus group housing and space allowance on responses of calves during open-field tests. Appl. Anim. Behav. Sci. 54, 109–121 (1997).

12. Valníčková, B., Stěhulová, I., Šárová, R. & Špinka, M. The effect of age at separation from the dam and presence of social companions on play behavior and weight gain in dairy calves. J. Dairy Sci. 98, 5545–5556 (2015).

13. Broom, D. M. & Leaver, J. D. Effects of group-rearing or partial isolation on later social-behavior of calves. Anim Behav 26, 1255–1263 (1978).

14. Meehan, C. L., Garner, J. P. & Mench, J. A. Isosexual pair housing improves the welfare of young Amazon parrots. Appl. Anim. Behav. Sci. 81, 73–88 (2003).

15. Glasper, E. R. & DeVries, A. C. Social structure influences effects of pair-housing on wound healing. Brain Behav. Immun. 19, 61–68 (2005).

16. Visser, E. K., Ellis, A. D. & Van Reenen, C. G. The effect of two different housing conditions on the welfare of young horses stabled for the first time. Appl. Anim. Behav. Sci. 114, 521–533 (2008).

17. Yarnell, K., Hall, C., Royle, C. & Walker, S. L. Domesticated horses differ in their behavioural and physiological responses to isolated and group housing. Physiol. Behav. 143, 51–57 (2015).

18. Baker, K. C. et al. Benefits of pair housing are consistent across a diverse population of rhesus macaques. Appl. Anim. Behav. Sci. 137, 148–156 (2012).

19. Grigg, E. K., Nibblett, B. M., Robinson, J. Q., Smits, J. E. Evaluating pair versus solitary housing in kennelled domestic dogs (*Canis familiaris*) using behaviour and hair cortisol: a pilot study. Vet Rec Open 4, e000193 (2017).

20. Costa, J. H. C., Meagher, R. K., von Keyserlingk, M. A. G. & Weary, D. M. Early pair housing increases solid feed intake and weight gains in dairy calves. J. Dairy Sci. 98, 6381–6386 (2015).

21. De Paula Vieira, A., von Keyserlingk, M. A. G. & Weary, D. M. Effects of pair versus single housing on performance and behavior of dairy calves before and after weaning from milk. J. Dairy Sci. 93, 3079–3085 (2010).

22. Gaillard, C., Meagher, R. K., von Keyserlingk, M. A. G. & Weary, D. M. Social housing improves dairy calves’ performance in two cognitive tests. PLoS One 9, e90205 (2014).

23. Meagher, R. K. et al. Effects of degree and timing of social housing on reversal learning and response to novel objects in dairy calves. PLoS One 10, e0132828 (2015).

24. Blanchette, I. & Richards, A. The influence of affect on higher level cognition: a review of research on interpretation, judgement, decision making and reasoning. Cogn Emot 24, 561–595 (2009).

25. Bradley, M. M., Cuthbert, B. N. & Lang, P. J. Picture media and emotion: effects of a sustained affective context. Psychophysiology 33, 662–670 (1996).

26. Roy, A. K. et al. Attention bias toward threat in pediatric anxiety disorders. J Am Acad Child Adolesc Psychiatry 47, 1189–1196 (2008).

27. Wright, W. F. & Bower, G. H. Mood effects on subjective probability asssessment. Organ Behav Hum Decis Process 52, 276–291 (1992).

28. Mendl, M., Burman, O. H. P., Parker, R. M. A. & Paul, E. S. Cognitive bias as an indicator of animal emotion and welfare: emerging evidence and underlying mechanisms. Appl. Anim. Behav. Sci. 118, 161–181 (2009).

29. Harding, E. J., Paul, E. S. & Mendl, M. Animal behavior: cognitive bias and affective state. Nature 427, 312 (2004).

30. Boleij, H. et al. A test to identify judgement bias in mice. Behav. Brain Res. 233, 45–54 (2012).

31. Mendl, M. et al. Dogs showing separation-related behaviour exhibit a ‘pessimistic’ cognitive bias. Curr. Biol. 20, 839–840 (2010).

32. Hintze, S., Roth, E., Bachmann, I. & Würbel, H. Toward a choice-based judgment bias task for horses. J Appl Anim Welf Sci 20, 123–136 (2017).

33. Düpjan, S., Ramp, C., Kanitz, E., Tuchscherer, A. & Puppe, B. A design for studies on cognitive bias in the domestic pig. J Vet Behav 8, 485–489 (2013).

34. Deakin, A., Browne, W. J., Hodge, J. J., Paul, E. S. & Mendl, M. A screen-peck task for investigating cognitive bias in laying hens. PLoS One 11, e0158222 (2016).

35. Baciadonna, L., Nawroth, C. & McElligott, A. G. Judgement bias in goats (Capra hircus): investigating the effects of human grooming. PeerJ. 4, e2485 (2016).

36. Neave, H. W., Daros, R. R., Costa, J. H. C., von Keyserlingk, M. A. G. & Weary, D. M. Pain and pessimism: dairy calves exhibit negative judgement bias following hot-iron disbudding. PLoS One 8, e80556 (2013).

37. Daros, R. R., Costa, J. H. C., von Keyserlingk, M. A. G., Hötzel, M. J. & Weary, D. M. Separation from the dam causes negative judgement bias in dairy calves. PLoS One 9, e98429 (2014).

38. Lecorps, B., Kappel, S., Weary, D. M. & von Keyserlingk, M. A. G. Dairy calves’ personality traits predict social proximity and response to an emotional challenge. Sci Rep 8, 16350 (2018).

39. Lalot, M., Ung, D., Péron, F., d’Ettorre, P. & Bovet, D. You know what? I’m happy. Cognitive bias is not related to personality but is induced by pair-housing in canaries (*Serinus canaria*). Behav. Processes 134, 70–77 (2017).

40. Hintze, S. et al. A cross-species judgement bias task: integrating active trial initiation into a spatial Go/No-go task. Sci Rep 8, 5104 (2018).

41. Papini, M. R. Comparative psychology of surprising nonreward. Brain Behav. Evol. 62, 83–95 (2003).

42. Roelofs, S. et al. Judgement bias in pigs is independent of performance in a spatial holeboard task and conditional discrimination learning. Anim Cogn 20, 739–753 (2017).

43. Gygax, L. The A to Z of statistics for testing cognitive judgement bias. Anim Behav 95, 59–69 (2014).

44. Hothorn, T., Bretz, F. & Westfall, P. Simultaneous inference in general parametric models. Biom J 50, 346–363 (2008).

45. Holson, R. & Sackett, G. P. Effects of isolation rearing on learning by mammals. Psychol Learn Motiv 18, 199–254 (1984).

46. Jones, S. et al. Assessing animal affect: an automated and self-initiated judgement bias task based on natural investigative behaviour. Sci Rep 8, 12400 (2018).

47. Bailoo, J. D. et al. Effects of cage enrichment on behavior, welfare and outcome variability in female mice. Front Behav Neurosci 12, 232 (2018).

48. Novak, J. et al. Effects of stereotypic behaviour and chronic mild stress on judgement bias in laboratory mice. Appl. Anim. Behav. Sci. 174, 162–172 (2016).

49. Holm, L., Jensen, M. B. & Jeppesen, L. L. Calves’ motivation for access to two different types of social contact measured by operant conditioning. Appl. Anim. Behav. Sci. 79, 175–194 (2002).

50. Cohen, S. & Wills, T. A. Stress, social support, and the buffering hypothesis. Psychol Bull 98, 310–357 (1985).

51. Rault, J. L. Friends with benefits: social support and its relevance for farm animal welfare. Appl. Anim. Behav. Sci. 136, 1–14 (2012).

52. Kiyokawa, Y., Hiroshima, S., Takeuchi, Y. & Mori, Y. Social buffering reduces male rats’ behavioral and corticosterone responses to a conditioned stimulus. Horm Behav 65, 114–118 (2014).

53. Færevik, G., Jensen, M. B. & Bøe, K. E. Dairy calves social preferences and the significance of a companion animal during separation from the group. Appl. Anim. Behav. Sci. 99, 205–221 (2006).

54. Duncan, I. J. H. Animal welfare defined in terms of feelings. Acta Agric. Scand. A Anim. Sci. 27, 29–35 (1996).

