## Supplementary material to Buckova et al_Manuscript for "Pair housing makes calves more optimistic"

### Details on the experimental procedure

**Sub-phases during the operant learning for trial initiation**

The operant learning for active trial initiation was divided into three sub-phases: (1) training calves to touch the trial initiator held next to the positive location (P) with the muzzle and to drink the reward (0.25 l of milk) given at P, (2) training calves to touch the trial initiator at a distance of 1.5 m from P and then to walk to P to get a reward, and (3) training calves to touch the trial initiator at a distance of 3 m from P and then to walk to P to get a reward.

In the first trial of the first session in **sub-phase 1** (following the habituation phase), a teat-bucket with milk was held close to the calf. As all calves were used to get milk from teat-buckets, they usually started to explore it during this trial. After the calf had finished drinking the milk, the bucket was removed. In the second trial, the bucket with the milk was held close to P, still within the arena. In the third trial, the trial initiator was introduced and from now onwards, the bucket was always placed at P from outside of the arena (placed in the bucket holder), so that calves could only see the red bucket teat. As soon as the calf touched the trial initiator, which was held very close to P in this sub-phase, it was lifted and then removed by the researcher. The bucket with the reward was placed into the bucket holder at P. After the calf had finished drinking, the bucket was removed and the trial initiator was made available again. The learning criterion was defined as making a maximum of two mistakes in two consecutive sessions. A mistake was defined as every interaction with the trial initiator that was not touching it with the muzzle. If a calf did not touch the trial initiator within 30 s or did not start drinking the milk within 30 s after presentation of the trial initiator, the session was terminated.

In **sub-phase 2**, the distance between the trial initiator held by the researcher and P was increased to 1.5 m. The learning criterion for this sub-phase defined as making a maximum of one mistake per session. A mistake was defined as not touching the trial initiator with the muzzle within 60 s after presentation. If a calf did not touch the trial initiator within 90 s or did not start drinking the milk within 60 s after having touched the trial initiator, the session was terminated.

In **sub-phase 3,** the distance between the trial initiator and P was increased to 3.0 m and the trial initiator was suspended from the ceiling and controlled by the researcher behind the wall with the five goal-holes via a pulley. The learning criterion for this sub-phase was defined as making a maximum of one mistake per session. A mistake was defined as not touching the trial initiator within 90 s after the calf had finished drinking the milk. If a calf did not touch the trial initiator within 120 s or did not start drinking the milk within 90 s after having touched the trial initiator, the session was terminated.

**Supplementary Table S1.**

**Number of excluded and additional test sessions per housing treatment.**

| Test session | Number of excluded test sessions | | Number of additional test sessions | |
|---|---|---|---|---|
| | IND | PAIR | IND | PAIR |
| 1 | 1 | 2 | 0 | 1 |
| 2 | 1 | 3 | 0 | 1 |
| 3 | 0 | 3 | 1 | 1 |
| 4 | 1 | 2 | 0 | 1 |
| Total | 3 | 10 | 1 | 4 |

Since a stable performance in positive and negative trials during test sessions is a prerequisite for a valid interpretation of the responses in ambiguous trials, only test sessions in which calves showed at least 13 Go responses in positive trials and 13 No-go responses in negative trials were included in the statistical analyses. The number of test sessions in which calves made more than four mistakes in positive trials and/or more than four mistakes in negative trials are shown here (excluded test sessions). Whenever possible within the 30 training days, calves were then tested in up to two additional test sessions.

**Supplementary Figures**

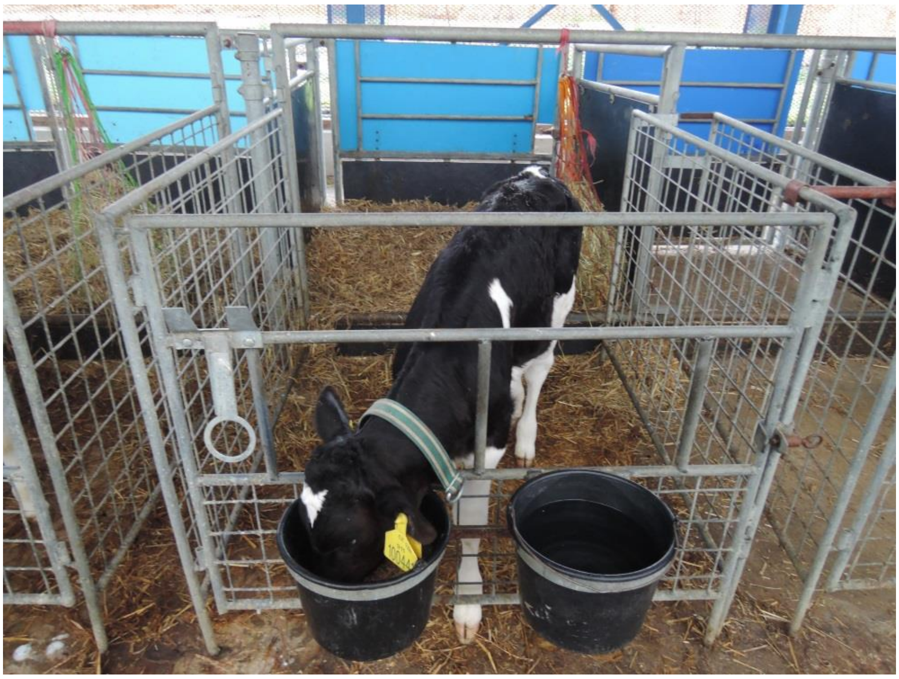

**Supplementary Figure S1.** An experimental calf housed individually.

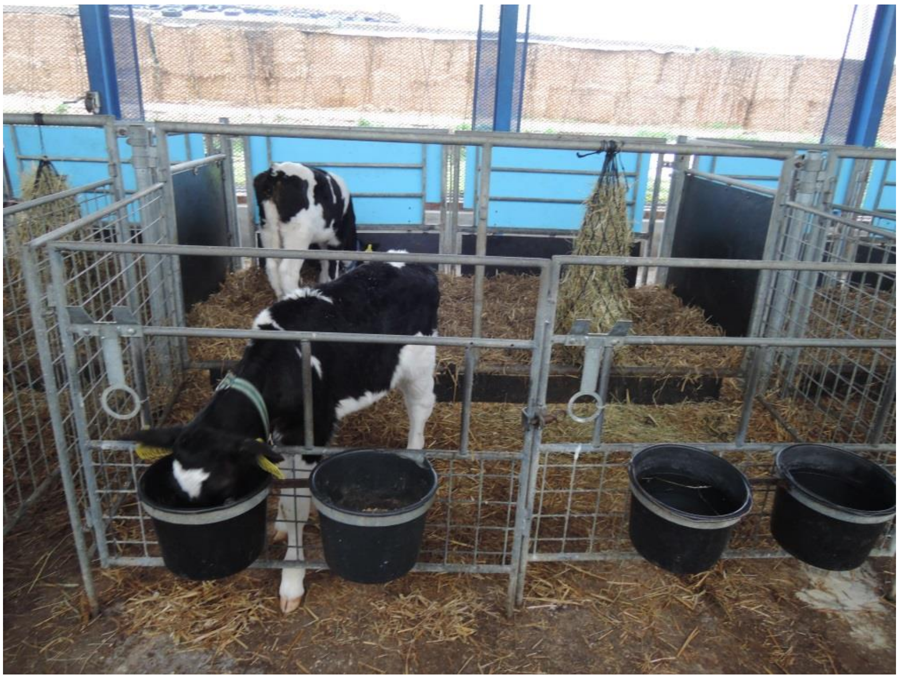

**Supplementary Figure S2.** Pair housing of the experimental calves.

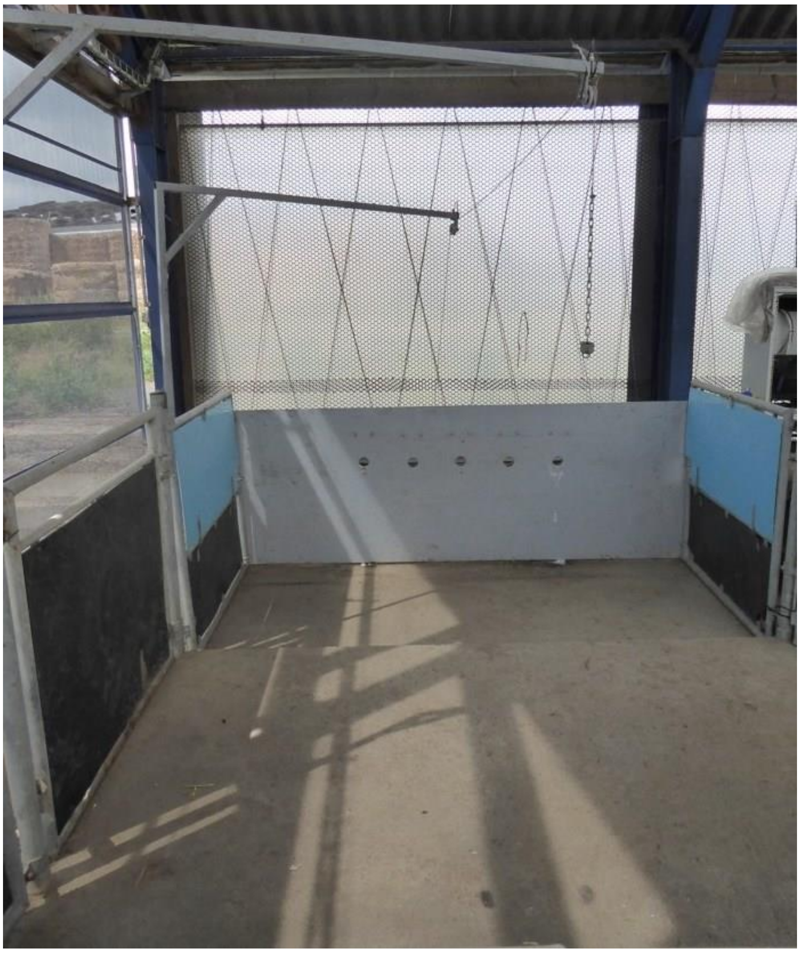

**Supplementary Figure S3.** Photograph of the experimental arena with a view on the wall with the five goal-holes where the teat of the bucket was presented during training/testing.

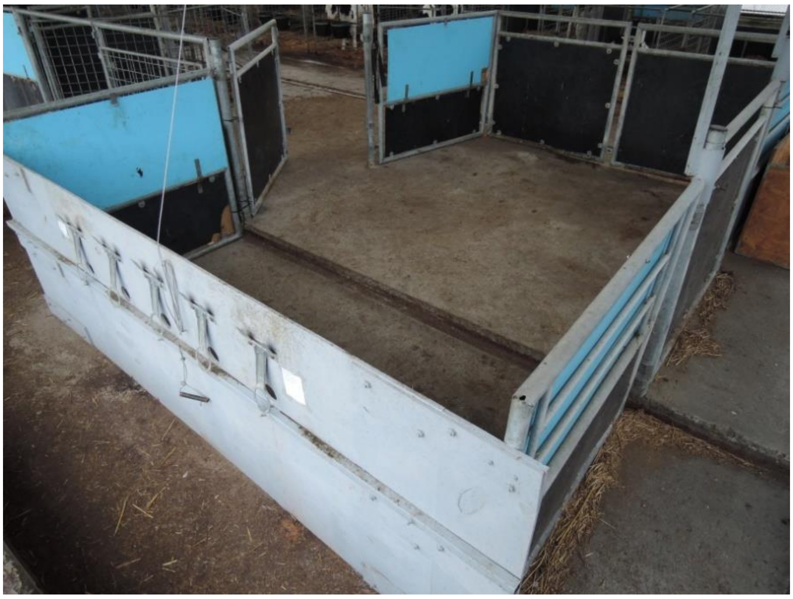

**Supplementary Figure S4.** Photograph of the experimental arena with a view on the wall with the five bucket holders where the teat-bucket was placed during training/testing.
